# Evaluation of High Arctic terrestrial habitats as potential hotspots of nitrous oxide emissions (Hornsund region, South Spitsbergen)

**DOI:** 10.64898/2026.02.27.708492

**Authors:** Julia Brzykcy, Renata Matlakowska, Jakub Grzesiak

## Abstract

Carbon dioxide (CO_2_), methane (CH_4_) and nitrous oxide (N_2_O) are the main greenhouse gases (GHGs) contributing to the ongoing climate crisis. Among those N_2_O has the highest global warming potential and is mainly of microbiological origin. Tropical rainforests are considered the primary natural source, although in recent years fluxes of N_2_O from polar ecosystems have been reported at comparable levels. In this study we aimed to identify High Arctic terrestrial habitats with the highest potential to become sources of N_2_O emissions. A microbiological and geochemical analysis was performed on soil procured from the biologically and geomorphologically diverse South Spitsbergen region in search of biotic and abiotic determinants of a N_2_O emission hotspot. Terrestrial sites within this High Arctic area vastly differed in their potential to emit substantial N_2_O amounts. External organic matter inputs were pivotal in maintaining a pool of inorganic nitrogen compounds for microbially-mediated N_2_O-generating processes such as denitrification. The examined planktivorous seabird colony presented a unique, potential N_2_O emission hotspot as it featured persistent acidification of the surrounding soil, a steady ammonia release and nitrate presence even after breeding season closure. Soils of the majority of analyzed sites did not display detectable nitrate and/or ammonia levels, with some areas having the characteristics of a N_2_O-sink rather than an emitter, especially postglacial moraine deposits. The presented data encourage further, more targeted investigations of High Arctic N_2_O emission hot- and coldspots to progressively improve N_2_O emission estimates for permafrost affected regions worldwide.

## 1. Introduction

Greenhouse gas (GHG) emissions due to fossil fuel combustion and agricultural practices have been repeatedly proven to be the cause of the ongoing climate change. Carbon dioxide (CO_2_), methane (CH_4_) and nitrous oxide (N_2_O) are the main GHGs contributing to global surface temperature rise. Among those, N_2_O has the highest global warming potential, being (per molecule) 300 times that of CO_2_ on a 100 years basis, and is now considered to be the dominant molecule contributing to the depletion of the stratospheric ozone layer with an atmospheric lifetime of 116 ± 9 years (Myhre et al., 2013; Tian et al., 2020). Global emissions of N_2_O originate predominantly from microbial processes, with negligible amounts accounting for abiotic sources – eg. chemodenitrification (Bothe et al., 2007). Tropical rainforests are considered the primary natural source, while anthropogenic sources mainly include agricultural soils subjected to fertilisation. The latter account for 43% of global N_2_O emissions and are responsible for atmospheric concentration increase from 280 to 334 ppb between 1920 and 2021 (Repo et al., 2009; Tian et al., 2020).

In recent years fluxes of N_2_O from some polar ecosystems have been reported at levels comparable to tropical forests and although emissions of this gas from polar and permafrost-affected soils are increasingly recognized as significant, they differ in scale from those in warmer and managed ecosystems (Repo et al., 2009). Tian et al. (2020) estimate that global agricultural soils are the dominant source, releasing up to 3.8 Tg NLO-N yr^−1^, with the whole agricultural sector (including pasture, manure management and aquaculture) totaling up to 5.8 Tg NLO-N yr^−1^. Natural soils also contribute substantially with values up to 6.5 Tg NLO-N yr^−1^ with tropical forest soils accounting for 2.2–3.7 Tg NLO-N yr^−1^, while wetlands and rivers act as additional hot spots (Tian et al. 2020; Zhou et al. 2021). Oceans can release as much as 4.3 Tg NLO-N yr^−1^ while natural inland waters, estuaries and coastal zones can contribute up to 0.4 Tg NLO-N yr^−1^. In contrast, Voigt et al. (2020) report that until recently, permafrost soils were considered negligible, yet upscaled N_2_O emission estimates for permafrost regions assume mean annual emissions of 1.27 Tg NLO-N yr^−1^. Despite this, polar emissions remain a small share of the global total, contributing roughly 7.1% of global emissions. However, due to their sensitivity to warming, permafrost thaw and hydrological changes it is suggested their relative importance may grow.

Several factors indicate that polar regions, especially those affected by permafrost are likely to become major contributors to the global N_2_O budget, significantly accelerating the current global warming event. Permafrost, which constitutes soils that remain at or below 0°C for at least two consecutive years, covers approximately 17% of the Earth’s land area – mostly in the Arctic, with a small percentage in Antarctica and alpine regions. It traps >50% of Earth’s organic carbon (C) stock and an unknown pool of N in the form of undecomposed animal and plant matter and microbial biomass (Jansson and Taş, 2014). Global temperature rise significantly contributes to the thawing of Arctic permafrost as air temperature in this region of the world has been rising >2 times faster than the global average in the past two decades (IPCC, 2019; Richter-Menge et al., 2017, Xie et al., 2022). Thawing of permafrost results in the release of substantial quantities of stored organic matter and mineral compounds into the active soil layer, accelerating the metabolic activity of soil microbial communities, particularly microorganisms involved in the decomposition of soil organic matter and the turnover of carbon and nitrogen, leading to the further emission of GHGs.

Since N_2_O has been proven to be a potent GHG and a stratospheric ozone depleting agent, efforts have been made to recognize and characterize its natural and man-made sources, pinpointing N_2_O emission hotspots (“patches that show disproportionately high reaction rates relative to the surrounding area”) and examining their features (McClain et al., 2003; Tian et al., 2020). The majority of this research focused on evaluating the N_2_O emitting potential of native and agricultural soils, while investigations of polar regions, majority of which concern the Arctic, concentrate on emissions of CO_2_ (resulting mostly from respiration) and CH_4_ (derived from archaeal methanogenesis and geogenic processes) leaving the processes responsible for N_2_O production poorly characterized, despite their significant impact on the global atmospheric budget (Convey et al., 2018; Voigt et al., 2020). Permafrost affected regions are by default very heterogeneous, as the majority of circumpolar terrestrial habitats fulfill these conditions. Investigated permafrost-based ecosystems with high fluxes included: thawing permafrost, permafrost peatlands, bare peat circles, ice-wedge polygons and seabird-affected taluses (Altshuler et al., 2019; Hayashi et al., 2018; Marushchak et al., 2021; Repo et al., 2009; Voigt et al., 2017, 2020). These studies, despite providing direct evidence of N_2_O emissions are rather one sided in terms of investigating the gasogenic components. While some of these authors take only the physicochemical properties of the soils into account, others overly focus on the microbial component, neglecting crucial abiotic variables. Furthermore, in the High Arctic the activity of only one feeding type of birds was investigated in connection to N_2_O releases, namely piscivores (fish eaters) and while their importance remains valid, there are other groups of marine animals that contribute labile nitrogen to the coastal permafrost soils which can stimulate microbial processes and subsequent emissions of gaseous N-compounds, including N_2_O (Zhu et al. 2012, Hayashi et al., 2018). Planktivorous birds (zooplankton-feeding) little auks (*Alle alle*) are known for their extensive colonies in that region where they deposit guano on a wide coastal area. However, despite numerous studies on their effects on the Arctic tundra, to date a direct relation to N_2_O emissions has not been made (Skrzypek et al., 2015; Zwolicki et al. 2013; Szymański et al., 2016).

As mentioned, N_2_O is mainly of microbiological origin with recognised several pathways leading to its emissions. Primary N_2_O production pathways involve direct mechanisms such as denitrification, which accounts for the majority of emissions (70% of global budget) (Mosier, 1998). In addition, although to a lesser extent, ammonia oxidation, nitrification, dissimilatory nitrate reduction to ammonium and the anammox process may serve as indirect sources of N_2_O, releasing it as a by-product (Kuypers et al., 2018). However, several prerequisites are needed to be met if a site is to become a N_2_O emission hotspot. High soil C and N with a low C:N ratio was proven to be a strong indicator of an N_2_O emitting site. Yet, whether a particular soil will act as a N_2_O source or a sink depends on a complex interaction between several biotic and abiotic factors (Wang et al., 2021). Soil moisture was considered as one of the main regulators of N_2_O emissions, that depends on a wide range of factors, including water content, water table level, bulk density, water holding capacity (WHC) and water-filled pore space (WFPS). Soil water content, besides nutrient dissolution and transport, impacts also O_2_ availability (Hu et al., 2015). Moderate water content of 60-70% was observed to promote development of both aerobic and anoxic niches in soil, which allows for ammonification, NH_4_^+^ oxidation and nitrification to take place, as well as anaerobic processes leading to N_2_O emissions derived from denitrification (Repo et al., 2009; Voigt et al., 2020). Nitrate content seems to play a pivotal and a very peculiar role in N_2_O release from soil environments. It is a direct substrate for denitrification, yet inhibits the reduction of N_2_O to N_2_ when present at high concentrations (Wang et al., 2021). Conditions, that promote N_2_O emissions also include low pH, which inhibits or delays the maturation of nitrous oxide reductase enzyme transcript leading to elevated N_2_O:N_2_ ratios (Liu et al., 2014). Vegetation cover is another factor influencing microbial N_2_O emissions as plants are known to compete with microorganism for resources, including the scarce in Arctic soils mineral N forms, like NO_3_^-^ and also organic N compounds, by which they inhibit processes leading to N_2_O fluxes (Ramm et al., 2022).

There are also microbial community related features that indicate potential N_2_O emission hotspots, such as presence of certain functional groups (eg. ammonia oxidizers, nitrifiers, denitrifiers), total cell abundance and activity, but also more specific markers like the ratio between the genes encoding nitrite reductase (*nirK*, *nirS*) and nitrous oxide reductase (*nosZ*), where higher nitrite reductase to nitrous oxide reductase gene ratios are typically indicative of increased soil N_2_O emissions (Hu et al., 2015; Voigt et al., 2020). Additionally, the bioavailability of iron, zinc and especially copper plays a role in the denitrification process, as those metals act as cofactors for several metalloenzymes involved in nitrate respiration pathways or accompanying reactions (Tavares et al., 2006).

In this study we aimed to identify High Arctic terrestrial habitats with the highest potential to become a source of N_2_O emissions. We procured soil samples from six sites within the diverse coastal landscape of the Hornsund fjord in South Spitsbergen and conducted an analysis of soil microbiological and environmental determinants including microbial community structure, abundance, diversity and metabolic versatility, as well as soil geochemical properties. We sought the answers to the following research questions: 1) How considerable are the differences in N turnover and N_2_O production potential between specific terrestrial sites within the Hornsund region; 2) Combinations of which biotic and abiotic factors (including external N inputs) are present in those soils that predispose them to become N_2_O emission sources or its sinks.

## 2. Materials and methods

### 2.1 Site description and sampling

Soil samples were collected in the vicinity of The Stanisław Siedlecki Polish Polar Station Hornsund located on Spitsbergen Island (Svalbard Archipelago, High Arctic). The Hornsund fjord area (South Spitsbergen) is characterised by diverse geomorphology and tundra vegetation types. Stations infrastructure is located 300 m from the shore of the Hornsund fjord, 1 km south of mountain ranges and 2.5 km west of the Hans Glacier (Wawrzyniak and Osuch, 2020). Fieldwork was conducted in summer of 2021 (25.08-01.09) during the 44th Institute of Geophysics Polish Academy of Sciences Polar Expedition and included six different microbial habitats (soil) (Fig. 1). Soil samples from each research site were collected in replicates (n=4) at a depth of 5-10 cm. Sampling points were located in the vertices of a square with each side equal to 100 cm. Samples were obtained with a spatula sterilised with 75% EtOH and flamed and then stored in Whirl-Pak polyethylene bags (Nasco, Madison, USA). Location was recorded using a GPSMAP® 276C (Garmin) device. Replicates from each sampling site were weighted, mixed, and divided into sub-samples at the stations laboratory and immediately frozen at – 20 °C until further processing in Warsaw, Poland.

**Fig. 1.**
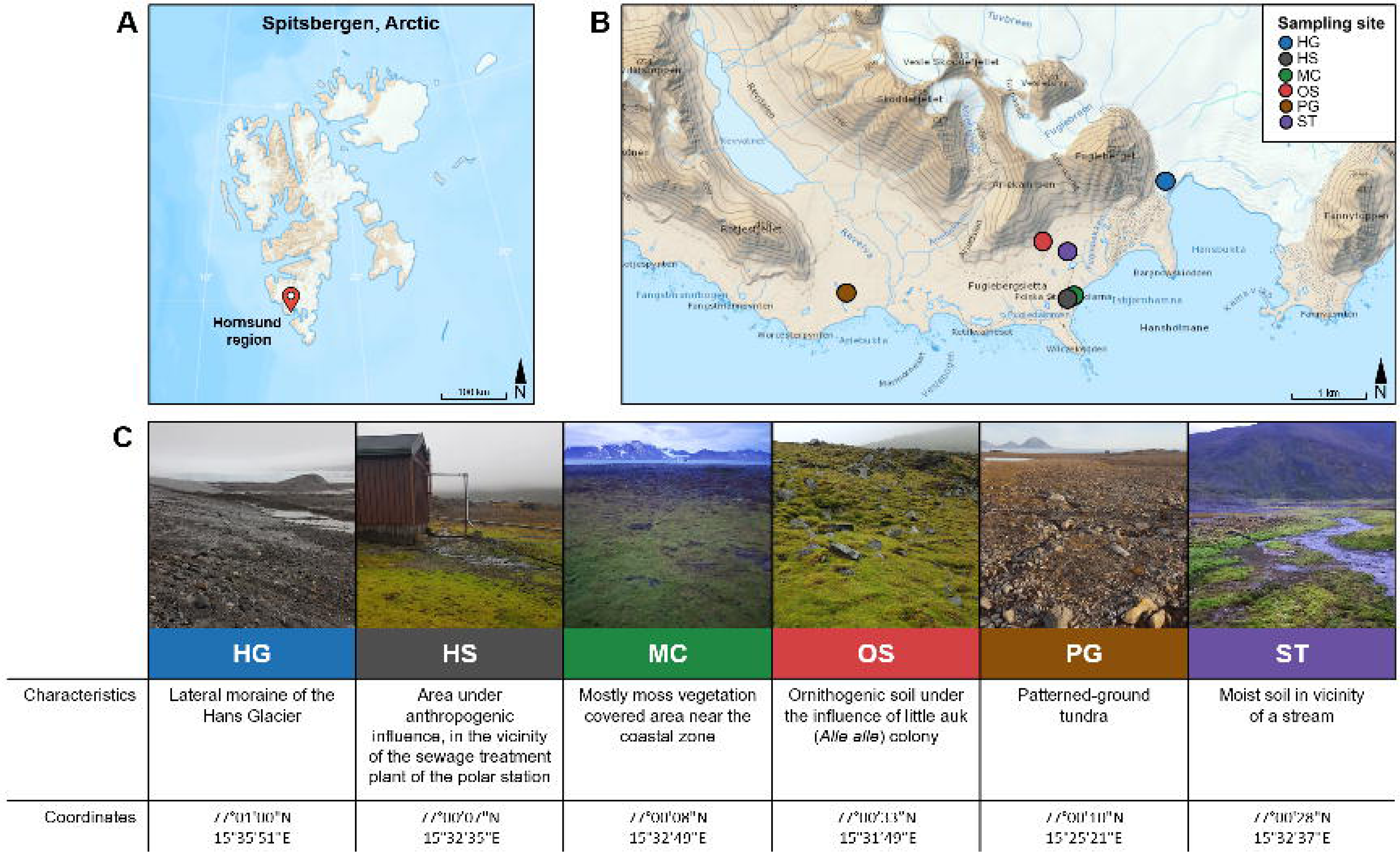
Localization of study sites. (A) Map showing Hornsund region localization on Spitsbergen map; (B) Map indicating locations of study sites; (C) Photos and characteristics of study sites. Maps were modified from TopoSvalbard (https://toposvalbard.npolar.no/).

### 2.2 Physicochemical analyses

Soil pH was measured using a 1:2 (wt:vol) soil suspension in 1M KCl according to standard methods using a SevenMulti pH meter with a InLab® 413 electrode (Mettler Toledo). Soil water (SW) content was determined after drying at 60 °C and soil organic matter (SOM) content was inferred after dry matter ignition at 500 °C. The total concentration of biogenic elements (CNS) was measured using the Flash 2000 apparatus (Thermo Fischer Scientific, Waltham, MA, USA). Total organic carbon was determined using Tyurin spectrophotometric method (FAO, 2021). Concentrations of NH_4_^+^, NO_3_^-^ and NO_2_^-^ were determined using NanoColor tests (Macherey-Nagel, Duren, Germany) according to the manufacturer’s instructions. The concentrations of selected metallic elements (Fe, Ca, Zn, Cu, Ni, Co, Pb) were determined by flame atomic absorption spectroscopy using SOLAAR M6 (TJA Solution, Cambridge, UK) with air and acetylene as a gas mixture following sample mineralisation in HNO_3_ (69%) in a laboratory microwave system (Ethos Plus, Milestone, Sorisole, Italy).

Volatile organic compounds were separated and identified as described further. One hundred grams of soil was placed in a serum bottle sealed with butyl rubber stoppers with aluminium caps and placed at 75 °C for 12 h. Using a gastight syringe, gases evolving during sample heating was collected from the tube. Analysis of collected gaseous samples was performed using gas chromatography (GC 7890A) coupled with mass spectrometry (MS 5973c) (Agilent Technologies, Santa Clara, CA, USA). PlotQ column was used (Agilent Technologies, Santa Clara, CA, USA). Five mililiter samples were taken using a gas-tight syringe, injected into the GC-MS and diluted in a helium stream 1:40 v/v (split). Separation of volatile compounds started at 70 °C, then the temperature was increased to 210 °C at a rate of 10 °C/minute. After reaching the temperature of 210 °C it was maintained for 3 minutes. The mass spectrometer scanned separated ions in the mass range from 10 to 300 Da, at an ionization of 70eV, filament temperature of 150 °C and 230 °C in the ionization chamber.

Non-volatile organic compounds were extracted from 20 g of soil using an automatic Soxhlet apparatus SER 158 (Velp Scientifica, Usmate Velate, Italy), with dichloromethane:methanol solution (8:2 v/v) for 4 h. The extract was dehydrated with anhydrous Na_2_SO_4_, and evaporated in a stream of nitrogen. Esterification was performed using BSTFA:TMCS (99:1 v/v) (Sigma Aldrich, Saint Louis, MO, USA) at 70 °C for 30 minutes. Esterified samples were stored at 4 °C for subsequent analysis. Separation of organic compounds in esterified samples was performed using gas chromatography (GC 7890A) coupled with mass spectrometry (MS 5973c) (Agilent Technologies, Santa Clara, CA, USA). Organic extracts were injected using an autosampler (7683 Series Injector) (Agilent Technologies, Santa Clara, CA, USA) in a volume of 5 µL. The standard deviation of injection according to the manufacturer’s data was a maximum of 0.3%. Injected samples were vaporised at 280 °C and diluted in a helium stream at a ratio of 1:5 v/v (split). Then, separation of organic compounds was performed using an HP-5MS column (30 m, 0.25 mm I.D., particle size 0.25 µm) (Agilent Technologies, Santa Clara, CA, USA), and helium as a carrier gas, at a flow of 1 mL/minute. Organic extracts were separated using a temperature gradient. For 5 minutes chromatographic column was heated at 100 °C, and then the temperature was increased to 280 °C, at a rate of 8 °C/minute. After reaching the temperature of 280 °C it was maintained for 20 minutes. The mass spectrometer scanned separated ions in the mass range from 40 to 800 Da at an ionization of 70eV, filament temperature of 150 °C and 230 °C in the ionization chamber. Chromatograms were integrated based on the degree of slope of the chromatogram baseline (threshold) and peak height. These coefficients were 15 and 0.01 respectively. The obtained mass spectra were identified using the Willey spectral library (version 3.2, Copyright 1988-2000 by Palisade Corporation with The Wiley Registry of Mass Spectral Data, 8th Edition with Structures, Copyright 2000 by John Wiley and Sons, Inc.) and NIST (version 2.0f The NIST Mass Spectral Search Program for the NIST/EPA/NIH). The database search parameters allowed for a 3% deviation of the analysed spectrum from the spectrum deposited in the library. Separation of organic compounds and calculation of their mass sums based on carbon chain length (C1-C5, C6-C10, C11-C15, C16-C20, C21-C30) were performed using a gas chromatograph (GC 7890A) equipped with a flame ionization detector (Agilent Technologies, Santa Clara, CA, USA). The injector port temperature was set at 300 °C, and the detector temperature at 320 °C. The flow rates for nitrogen, air, and hydrogen were 25 mL/min, 300 mL/min and 40 mL/min, respectively. Samples (volumes 5 µL) were injected with a split ratio of 1:5 into the HP-5MS column (column parameters: 30 m, 0.25 µm I.D., 0,25 µm film thickness, Agilent Technologies, Santa Clara, CA, USA). Helium was used as the carrier gas with an injection speed of 1 mL/min. Samples were then separated using a temperature gradient (four-step program: (i) 5 min, 40 °C; (ii) 10 min, temperature increase to 60 °C at a rate of 4 °C/min; (iii) 5 min, temperature increase to 100 °C, at a rate of 4 °C/min; (iv) 10 min, temperature increase to 300 °C, at a rate of 8 °C/min. Acetone, hexane, eicosane, and octadecanoic acid were used as standards in the applied calculations method.

### 2.3 Microbiological analyses

Microbial cells were extracted from soil following the procedure in Znój et al. (2022). One g of soil was added to 20 mL of a solution containing 0.9% (w/v) NaCl and 10 mM tetrasodium pyrophosphate (Na_4_P_2_O_7_) and shaken at 1000 rpm for 30 minutes at 4 °C. Then the sample was sonicated in a 4 °C water bath for 5 minutes, vortexed and centrifuged at 129 rcf for 1 minute to remove residual soil and recover cells suspended in the supernatant. The resulting suspension was used for the further described analyses.

Total prokaryotic cell count was determined by epifluorescence microscopy. Briefly, 1 mL of cell suspension was mixed with 90 μL of 25% glutaraldehyde (2% final conc.) and stained with SYBR Green I (2.5%) for 15 minutes in the dark (Lunau et al., 2005). Samples were filtered down through a NucleoporeTM black polycarbonate 0.2 μm pore size membranes (Whatman^®^) and air dried in the dark. Microbial cells were counted under the Nikon E-200 microscope with a 100 W Hg lamp and 100 × CFI 60 oil immersion objective, with a digital DS-Fi3 high-definition color microscope camera equipped with a 5.9 megapixel CMOS image sensor, and a filter block of wavelengths EX 450-490, DM 505, BA 520. Images of fields were analyzed in a Nikon NIS Elements BR 2.30 and MultiScan v. 14.02 computer scanning system. A minimum of 400 cells in 20 fields per sample were counted automatically in the image analysis system. The total microbial cell count was then evaluated through image analysis (Sieracki et al., 1985; Świątecki, 1997).

Community-Level Physiological Profiling was performed using the Biolog EcoPlate assay, which contains 31 carbon sources (in triplicates) on a 96-well plate, amended with a tetrazolium redox dye for indication of microbial respiration. Isolated microbial community in the form of cell suspension was added to each well in 100 μL aliquots. Plates were incubated for 6 weeks at 5°C, in the dark. Then, colour development was measured on the Varioskan™ LUX multimode microplate reader (Thermo Scientific™) at 590 nm wavelength in triplicates. Data were blanked against the T0 reading and then again, against the control well without a carbon source.

DNA was extracted using a Genomic Mini AX Soil Spin kit (A&A Biotechnology, Gdańsk, Poland), according to the manufacturer’s instructions. DNA concentration and quality determination were performed using an NanoPhotometer NP80 (Implen, München, Germany). The amplification of 16S rRNA gene V3-V4 region was performed using in triplicate using the WALK polymerase (*Pwo* polymerase with proof-reading activity) and xtV3F and xtV4R primers containing the Nextera XT adapter sequences, compatible with Illumina index and sequencing adapters. PCR products purification was performed using a Clean-Up kit (A&A Biotechnology, Gdansk, Poland), according to manufacturer instructions. For each sampling site, PCR reaction products were pooled together and sequenced on Illumina MiSeq apparatus (Illumina Inc., San Diego, CA, USA) in the DNA Sequencing and Synthesis Facility, Institute of Biochemistry and Biophysics, Polish Academy of Sciences (Warsaw, Poland). Sequencing was performed in paired-end mode (2 x 200 bp) and v3 (600 cycles) chemistry cartridge was used, which resulted in generations of long paired reads covering amplicons.

### 2.4 Data analysis

Microbiome taxonomic profiling was performed with help of the EzBioCloud platform using the PKSSU4.0 database (Yoon et al., 2017). Raw sequencing data were cleaned, aligned, and classified automatically. Chimeric, low quality, and non-target amplicons (chloroplast, mitochondrial, and archaeal) were automatically discarded. Illumina reads were deposited in the NCBI Sequence Read Archive (SRA) as BioProject PRJNA1068972. PICRUSt (Langille et al., 2013) was used to predict N cycling functionality using the metagenome_contributions.py script and the -l option to specify KEGG orthologs in the output for N fixation (K02588, K02586, K02591, K00531), nitrification (K10944, K10945, K10946, K10535), denitrification (K00368, K15864, K04561, K02305, K00376), dissimilatory nitrate reduction (K00370, K00371, K00374, K00373, K02567, K02568, K00362, K00363, K03385, K15876), assimilatory nitrate reduction (K00367, K10534, K00372, K00360, K00366, K17877), nitrate/nitrite transport (K02575), and ammonia assimilation (K00264, K00265, K00266, K01915, K01948). Relative abundance values were obtained as predicted gene copy numer to 16S rRNA copy number quotient.

All results were compiled using Excel 2016 (MS Office) for Windows. One-Way ANOVA and Tukey’s test were used to asses statistical significance. Correlations between family-rank sequence abundances derived from amplicon data and other environmental data were calculated using Pearson’s correlation coefficient. Principal Component Analysis (PCA) was performed using the singular value decomposition method. Data visualization and statistical analysis have been performed using the R software (R v.4.0.2) and the following packages: ggplot2, pheatmap, fmsb, Hmisc, ggpubr, multcompView, corrplot, and ggfortify (R Core Team 2002).

## 3. Results

### 3.1 Study site designations and features

Six study sites were chosen to capture contrasts in topology, geomorphology, and tundra vegetation, which in turn influenced their geochemical and microbial characteristics (Fig. 1). Acronyms of those sites are used in subsequent sections. The Hans Glacier (HG) moraine, devoid of vegetation and covered by loose gravel over ice, represented the most extreme and barren conditions. The Hornsund Station (HS) site, situated near the Stations sewage treatment facility, was sparsely colonized by *Sanionia uncinata* moss and a thin cyanobacterial mat. The Moss Coast (MC) formed a permanently wet, moss-dominated carpet, while the Ornithogenic Soil (OS) site beneath a little auk (*Alle alle*) colony supported dense, species-rich vegetation. Patterned Ground (PG) combined fine mineral material and rock fragments with scattered lichens and *Saxifraga* spp., whereas the Stream (ST) site comprised saturated moss “carpets.” The six sites represent a strong gradient from barren glacial forefield (HG) to biologically enriched ornithogenic soil (OS), providing a natural framework for comparing how vegetation and geomorphology shape soil properties. These diverse environments provided the physical and biological context for subsequent geochemical, organic, and microbial differences.

### 3.2 Geochemical environmental background

Soil chemistry reflected the contrasts in site conditions. Soil pH differed significantly, ranging from strongly alkaline HG soils (av. 9.5) to acidic OS soils (av. 5.5). PG (av. 8.2) and HS (av. 7.9) were moderately alkaline, whereas MC (7.6) and ST (7.1) were closer to neutral (Fig. 2A). Water content was especially high in ST (57.3%) and OS (33.9%), while other sites ranged from 10.7–19.3% (Fig. 2B). Soil organic matter (SOM) was greatest in MC (14.2%) and lowest in HG (1.6%) (Fig. 2C). Total carbon concentrations peaked at OS (170 g kgL¹ dm) but did not differ significantly from ST (156.4 g kgL¹ dm), MC (124.7 g kgL¹ dm), and HG (105.4 g kgL¹ dm) (Fig. 2D). Total nitrogen was highest in OS (48.0 g kgL¹ dm) and ST (41.3 g kgL¹ dm), while HG contained only 9.8 g kgL¹ dm (Fig. 2E). Sulfur showed a similar pattern, with OS and ST highest (35.3 and 32.5 g kgL¹ dm, respectively) (Fig. 2F). Organic carbon varied widely, from MC (69.5 g kgL¹ dm) to HG (6.4 g kgL¹ dm), with intermediate values at ST (37.5 g kgL¹ dm) and OS (23.7 g kgL¹ dm) (Fig. 2G). C:N ratios were greatest in nutrient-poor HG (10.8) and PG (6.5), but lowest in nutrient-enriched OS (3.5) and ST (3.8) (Fig. 2H).

**Fig. 2.**
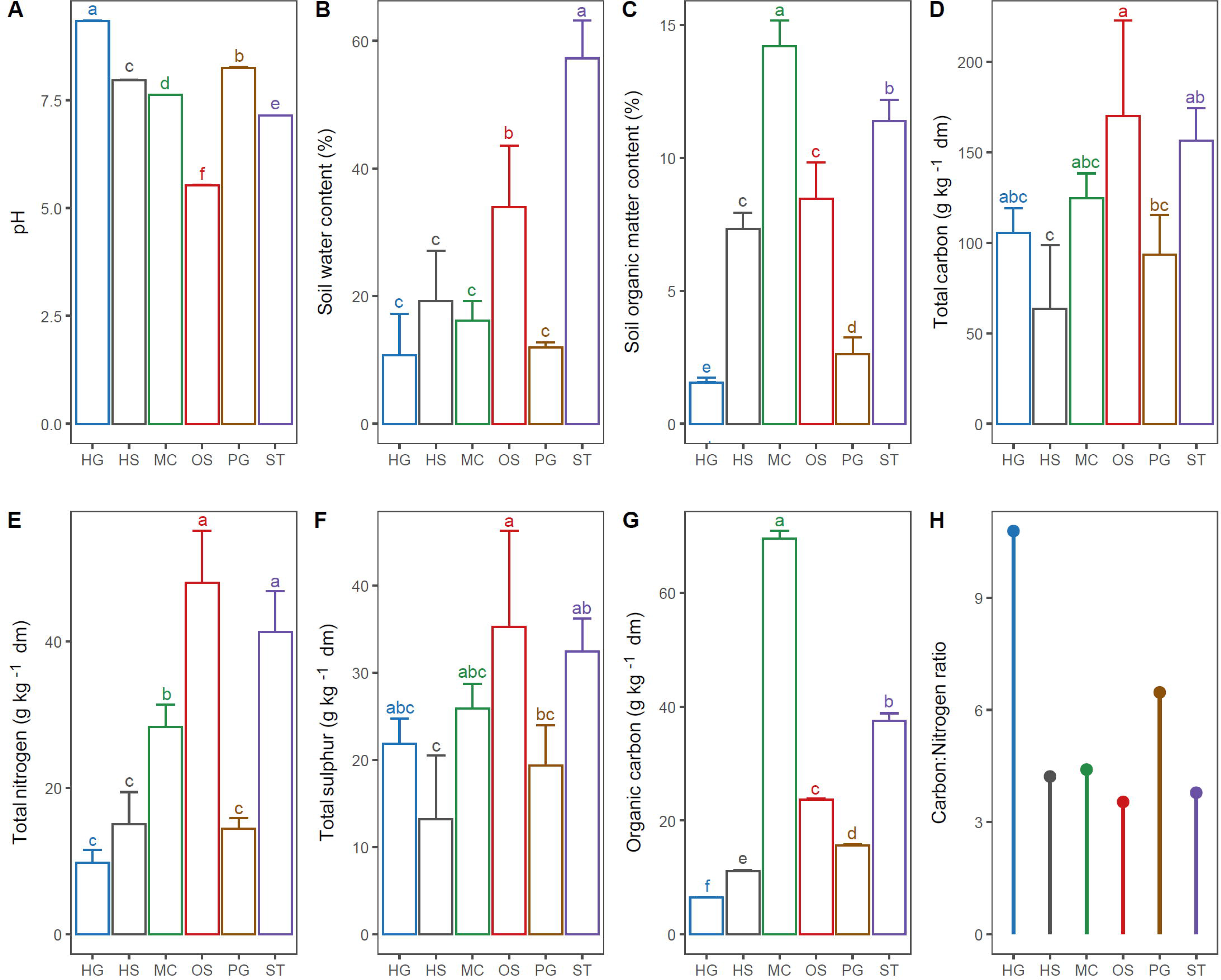
Basic geochemical parameters of examined High Arctic soils. A) Soil pH; B) Soil water content (% fresh mass); C) Soil organic matter content (% fresh mass); D) Total carbon concentration (g kg^-1^ dm); E) Total nitrogen concentration (g kg^-1^ dm); F) Total sulphur concentration (g kg^-1^ dm); G) Organic carbon concentration (g kg^-1^ dm); H) Total carbon to total nitrogen ratio; dm – dry mass. Values represented by bars with different letters differ significantly (*p* adj <0.05).

Trace element profiles underscored these contrasts (Table 1). ST and OS were Fe-rich (77.41 and 52.87 g kgL¹ dm), whereas Ca dominated HG and PG (105.24 and 43.74 g kgL¹ dm). Zn exceeded 200 mg kgL¹ dm in MC, HS, and ST, while Cu was elevated in ST and OS (53.63 and 20.65 mg kgL¹ dm, respectively). Ammonium was most abundant in ST (28.09 mg kgL¹ dm), followed by OS (18.16 mg kgL¹ dm) and MC (9.54 mg kgL¹ dm). Nitrate occurred only in OS (3.33 mg kgL¹ dm), whereas nitrite was detected at low levels across all sites (0.04–0.22 mg kgL¹ dm).

**Table 1.**
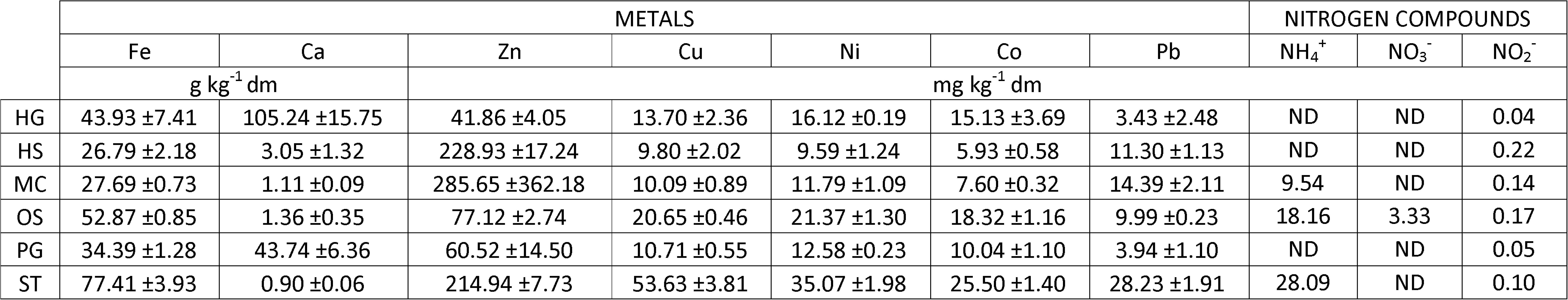
Concentration of metals and inorganic nitrogen compounds in examined High Arctic soils; dm – dry mass.

In summary, sites rich in vegetation and water (OS, ST, MC) contained higher nutrient and organic matter levels, while barren HG soils were nutrient-poor, alkaline, and Ca-dominated.

### 3.3 Organic compound composition

Organic compound profiles reflected the geochemical gradients. Aliphatic fractions (C1–C30) showed similar class contributions across sites (C1–C5: 15.7±1%; C6–C10: 31.5±5%; C11–C15: 36±2%; C16–C20: 16.5±1.8%; C21–C30: 0.3±0.2%) but varied in total abundance, peaking in ST (103.9 g kgL¹ dm) and lowest in HS (24.9 g kgL¹ dm) (Fig. 3A). Volatile compounds released by heating displayed site-specific signatures: 1-butanol dominated OS (10.3%), trichloromethane dominated HS (3.9%), PG (3.4%), MC (3.0%), and HG (2.02%), whereas trifluoroacetamid dominated ST (3.8%). Ethanol was broadly released across sites, most strongly in ST, MC, and HS (2.5–2.6%). Distinct patterns also emerged in compound families (Fig. 3C): HG soils were enriched in alkenes, acids, and sugars, MC in alkanes, ethers, and phenolics, and ST in alcohols, sterols, and N-related compounds. OS soils reflected mixed inputs, consistent with bird guano enrichment.

**Fig. 3.**
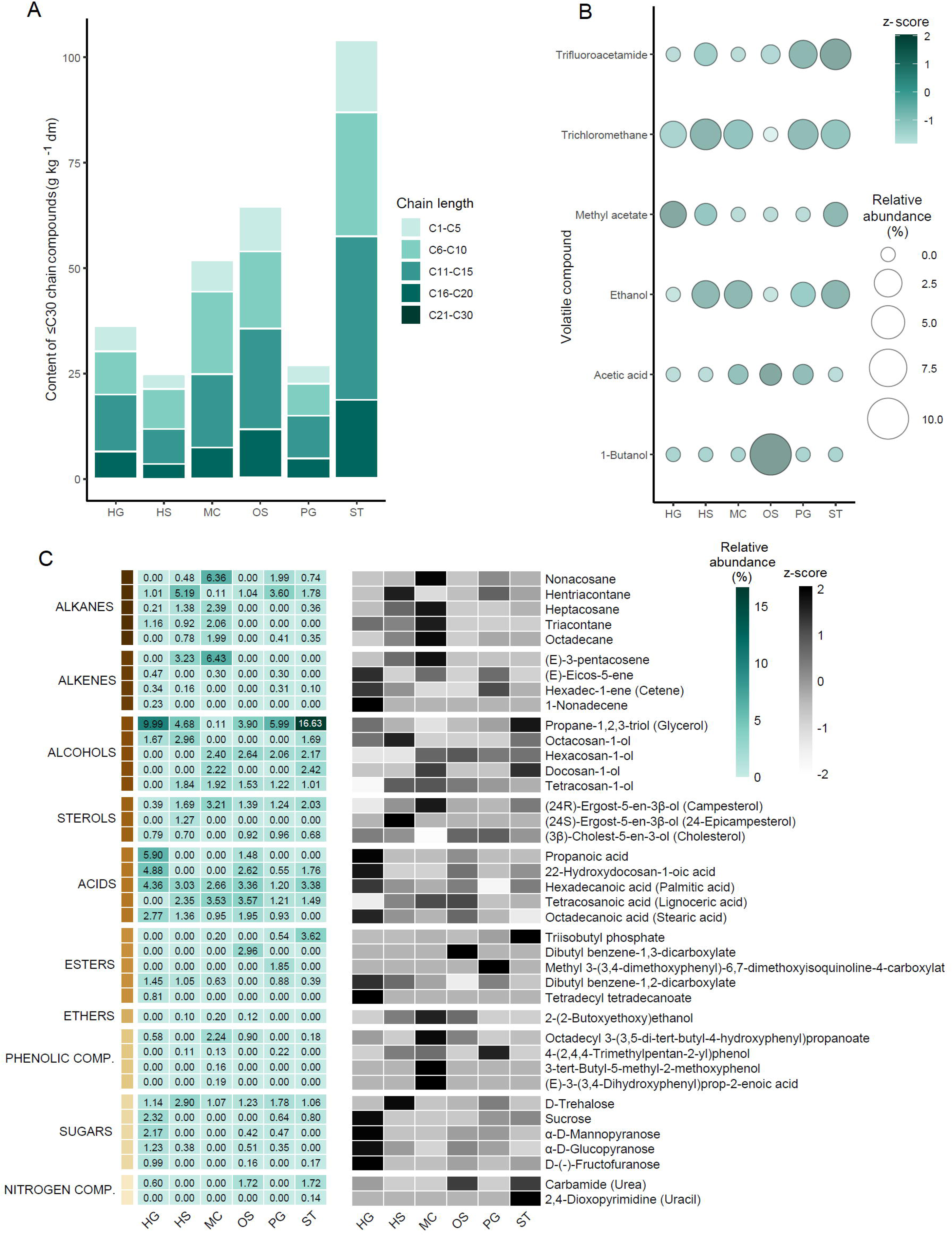
Quantity and quality of organic compounds in examined High Arctic soils. A) Concentration of aliphatic (≤30C) chain-containing compounds (g kg^-1^ dm) and their percentage contribution (by mass) by chain length; B) Relative abundance of major organic volatiles released by soil samples; C) Relative abundance of recognized soil organic compounds – percentage contribution (right) and row-scaling z-score (left); dm – dry mass.

In summary, organic compound pools showed clear site “fingerprints,” with barren HG soils dominated by simple compounds, bird-influenced OS soils enriched in alcohols and N-compounds, and waterlogged ST soils supporting the most diverse chemical profiles.

### 3.4 Microbial community abundance and diversity

Microbial parameters mirrored mentioned environmental contrasts. Prokaryotic cell counts spanned four orders of magnitude, from OS (7.39 × 10¹L ± 2.35 × 10¹L cells gL¹ dm) to HG (2.60 × 10L ± 2.45 × 10L cells gL¹ dm), with intermediate values in ST (3.53 × 10L ± 3.32 × 10L), MC (1.42 × 10L ± 5.46 × 10L), HS (6.80 × 10L ± 4.41 × 10L) and PG (2.44 × 10L ± 1.53 × 10L) (Fig. 4A). Diversity followed a similar trend: Shannon indices ranged from 7.21 in MC and ∼7.0 in HS/OS to just 3.18 in HG (Fig. 4B). Metabolic diversity was highest in ST (3.26), MC (3.25), and OS (3.23), lower in HS and PG (2.74), and lowest in HG (2.06) (Fig. 4C).

**Fig. 4.**
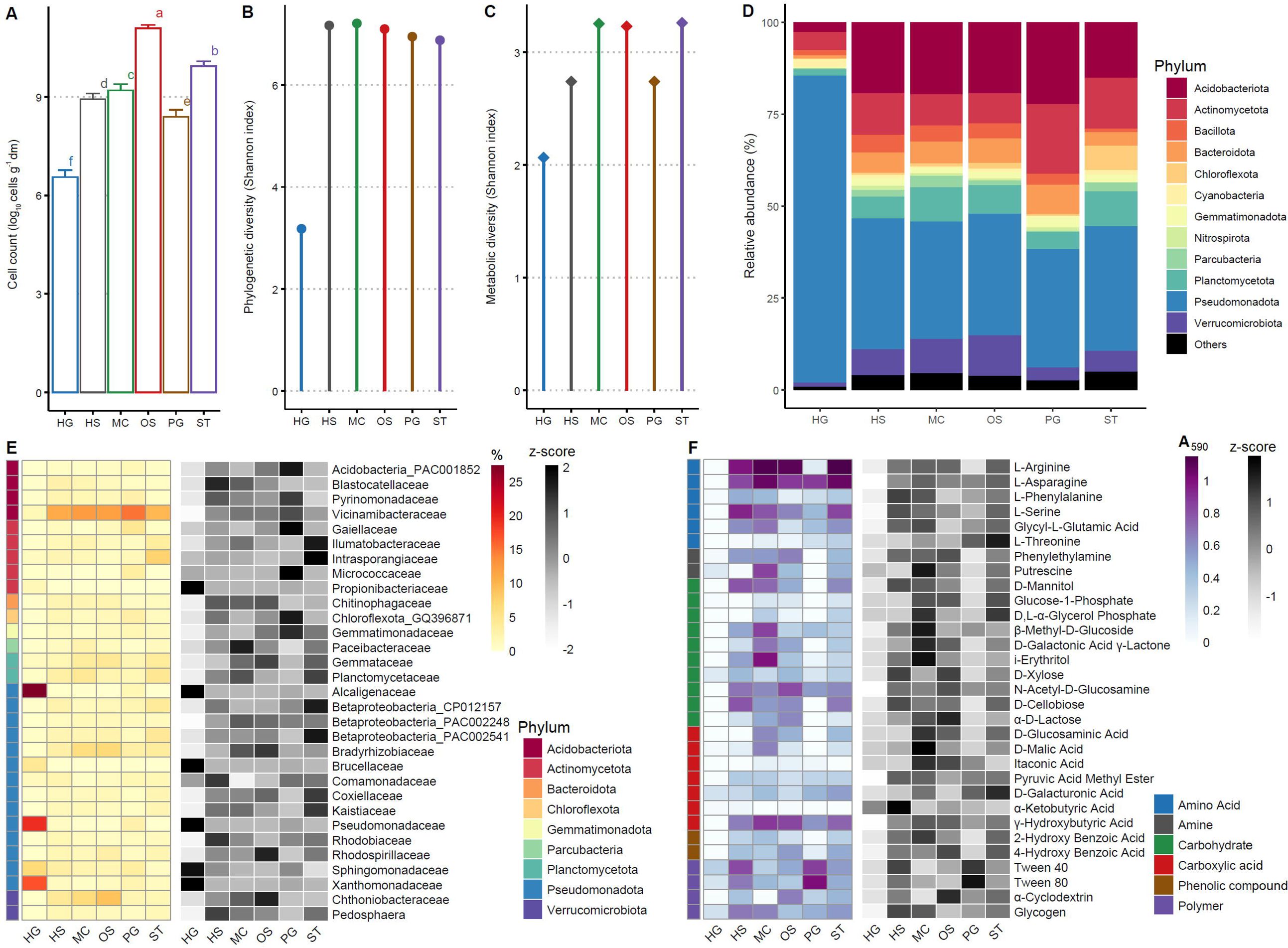
Microbiological parameters of examined High Arctic soils. A) Prokaryotic cell count; B) Phylogenetic diversity – Shannon index α diversity based on 16S rRNA gene amplicon datasets; C) Metabolic diversity - Shannon index α diversity based on Biolog EcoPlate responses; D) Relative abundance of phylum-ranked sequences; E) Relative abundance of family-ranked sequences – percentage contribution (right) and row-scaling z-score (left); F) Community responses on Biolog EcoPlates – mean A_590_ values from 3 replicates (right) and row-scaling z-score (left).

Bacterial community phylogenetic composition also reflected site conditions (Fig. 4D). All sites were dominated by Pseudomonadota, especially HG (83.6%), which had a uniquely skewed profile compared to the more balanced communities of other sites (32–35.5%). Acidobacteriota were common in PG (22.34%), OS (19.39%), MC (19.64%), HS (19.41%), and ST (15.07%), but rare in HG (2.61%). Actinomycetota peaked in PG (18.86%) and ST (13.96%). Verrucomicrobiota were enriched in OS (11%) and MC (9.3%). Family-level patterns confirmed site-specific associations (Fig. 4E): *Vicinamibacteraceae* dominated most sites (11.9±2%) but were nearly absent in HG, where *Alcaligenaceae* (27.3%), *Pseudomonadaceae* (19.2%), and *Xanthomonadaceae* (16.9%) prevailed. OS soils were enriched in *Chthoniobacteraceae*, PG in *Gaiellaceae*, ST in *Intrasporangiaceae*, and *Bradyrhizobiaceae* were broadly present (2.8–6.7%) except in HG.

Biolog EcoPlate assays reinforced these differences (Fig. 4F). Non-glacial soils (OS, ST, MC, HS, PG) displayed strong utilization of amino acids, carbohydrate derivatives, and polymers, whereas HG showed weak responses, underscoring the impact of nutrient scarcity and extreme conditions.

In summary: richly vegetated OS, ST, and MC soils supported abundant, diverse, and metabolically versatile microbial communities, while HG soils hosted sparse, low-diversity assemblages dominated by stress-tolerant taxa.

### 3.5 Functional gene potential for nitrogen cycling

Predicted functional profiles (PICRUSt) showed widespread capacity for nitrogen cycling (Fig. 5). Genes for NL fixation, nitrification, denitrification and nitrate reduction were present across sites, with denitrification and dissimilatory nitrate reduction genes (nirB, nirD, nirK) being most abundant. Relative contributions of nitrogen cycling genes were highest in HG (57%), though the observed low cell density meant absolute gene copy numbers were minimal. When corrected for cell abundance, OS soils harbored the largest pool of N-cycling genes, followed by ST. Ratios of nitrite reductase to nitrous oxide reductase (nirK+nirS/nosZ) highlighted potential NLO emission hotspots, particularly ST (3.1) and HS (2.5), compared with OS (1.3) and HG (1.1).

**Fig. 5.**
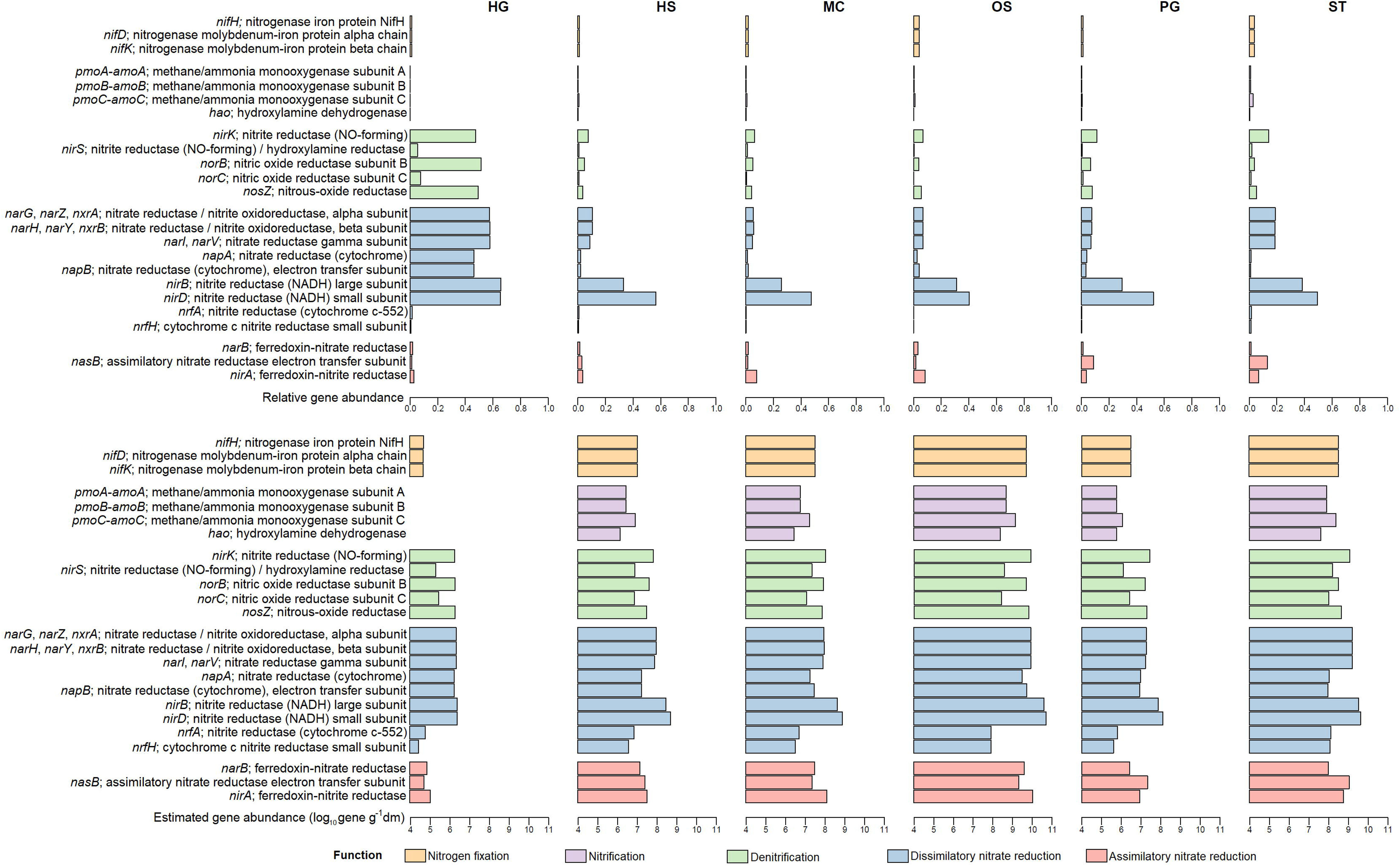
Presence of nitrogen-cycling genes in examined High Arctic soils as predicted by PICRUSt: relative abundance (upper graph), estimated abundance per g soil dm (lower graph); dm – dry mass.

In summary, although HG soils were enriched in N-cycling genes in relative terms, only nutrient-rich OS and ST soils combined high gene abundance with large microbial populations, making them the most functionally significant for nitrogen cycling.

### 3.6 Linking community composition with NLJO emission drivers

Correlation analyses integrated the biotic and abiotic datasets (Fig. 6A). C:N ratio negatively correlated with families such as *Chitinophagaceae*, *Coxiellaceae*, *Kaistiaceae* and *Betaproteobacteria*_PAC002248. By contrast, *Gemmataceae* and *Coxiellaceae* correlated positively with cell abundance, metabolic diversity, and total N and C. Principal Coordinate Analysis (Fig. 6B) placed OS within close proximity of ST as two nutrient- and biomass-rich hotspots, characterized by high microbial diversity and strong functional potential. Key drivers included C:N ratio, water content, and pH, with PC1 and PC2 explaining 77.57% of total variance.

**Fig. 6.**
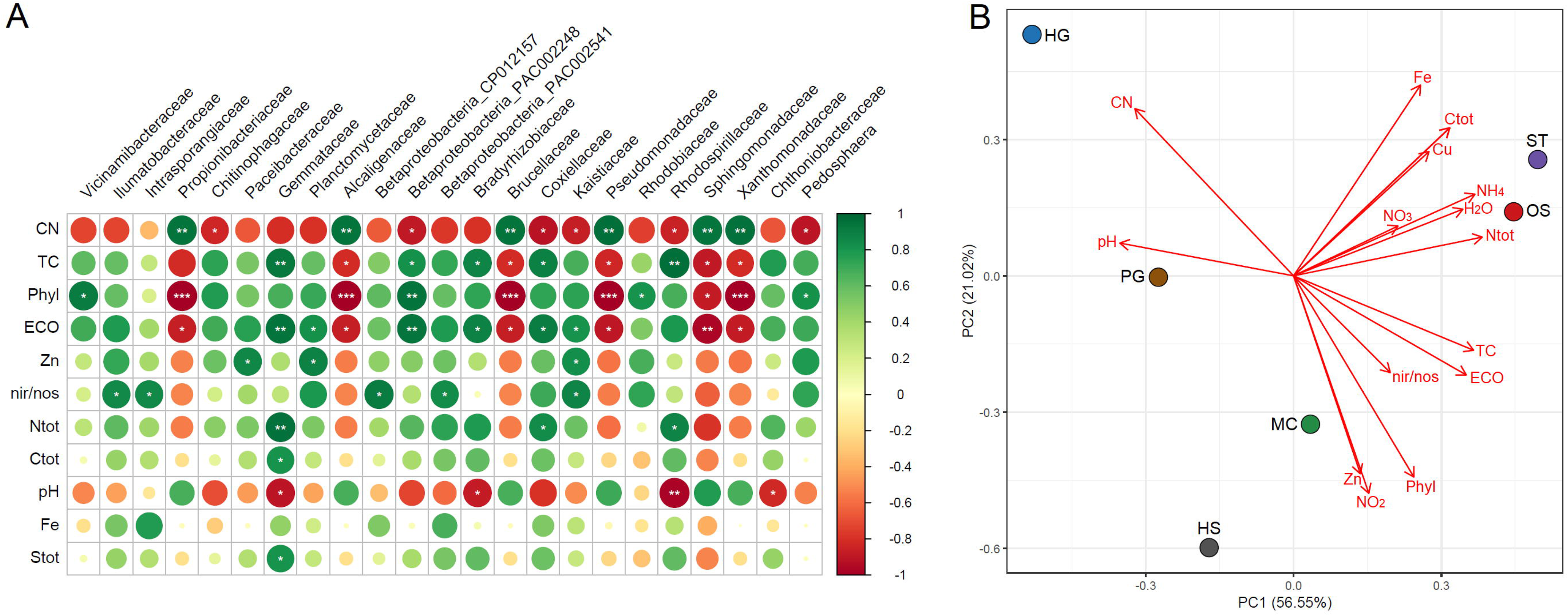
A) Correlogram of soil family-rank bacterial sequence abundance, soil physicochemical parameters and the relative (%) and estimated (log_10_ g^-1^ dm) abundance of the nitric oxide reductase subunit C gene *NorC*. Asterisks represent statistical significance levels: * p<0.05, ** p<0.01, *** p<0.001). CN - total carbon to total nitrogen ratio; TC - prokaryotic cell count; Phyl - phylogenetic diversity (Shannon index α diversity based on 16S rRNA gene amplicon datasets); B) Principal component analysis (PCA) of selected geochemical and biological data. ECO - Metabolic diversity (Shannon index α diversity based on Biolog EcoPlate responses); NorC - estimated (log_10_ g^-1^ dm) abundance of the nitric oxide reductase subunit C gene *NorC*; NorC% - relative (%) abundance of the nitric oxide reductase subunit C gene *NorC*; Ntot - Total nitrogen concentration (g kg^-1^ dm); Ctot - Total carbon concentration (g kg^-1^ dm); Fe -Total iron concentration (g kg^-1^ dm); Stot - Total sulphur concentration (g kg^-1^ dm), NO_3_^-^ - nitrate concentration (mg kg^-1^ dm), NO ^-^ - nitrite concentration (mg kg^-1^ dm); NH_4_^+^ - ammonia concentration (mg kg^-1^ dm), H_2_O – water content (% weight in fresh mass).

Consequently, nutrient availability, soil moisture, and stoichiometry (C:N) jointly shaped microbial communities and their role in NLO emission, with OS and ST emerging as the most active hotspots.

## 4. Discussion

### 4.1 Breeding colony of marine, planktivorous birds of the species little auk (Alle alle) is the most probable N_2_O emission hotspot

In our analysis two sites emerged as likely hotspots of N_2_O emissions, namely ST and OS. Especially the later showed traits characteristic of a potential source of microbially-produced nitrous oxide, mainly due to the high total nitrogen concentration, detectable nitrate amounts and an abundant and diverse microbial community. This site contains a breeding colony of marine, planktivorous birds of the little auk species which is responsible for the formation of the surrounding lush nitrocoprophilous tundra (Szymański et al., 2016). Piscivorous seabird activity has been recognized as a significant factor in boosting N_2_O emissions in Arctic tundra ecosystems, however plankton-feeders’ effect was not examined to date (Zhu et al., 2012; Hayashi et al., 2018). This colony site was investigated in terms of nitrogen compound concentration before by Zwolicki et al. (2013), where NO_3_^-^ concentration as high as 350 mg kg^-1^ were recorded and concentrations of NH_4_^+^ at 26 mg kg^-1^. While ammonia concentrations were comparable, the NO_3_^-^ concentrations were much higher than those recorded in our study. The time of sample collection might be the cause of these discrepancies as late August/early September is far past the birds’ peak activity (Ribeiro et al., 2024). This would indicate, that the site experiences “hot moments” of N_2_O emissions as nitrous oxide fluxes were found to significantly correlate with nitrate content in sea bird affected areas (Hayashi et al., 2018). However, our data indicate a possibility of low-rate N_2_O emissions even past the breeding season due to the relatively high ammonia concentrations still present at that time. This might be an exclusively planktivorous bird colony-associated phenomenon. Compared to piscivorous birds, the guano of planktivors contains (beside proteins and uric acid) a substantial amount of chitin, which is more resistant to degradation and consequently to ammonification then other guano components, hinting towards a gradual decomposition process and a constant ammonia release (Szymański et al., 2016; Jacquiod et al., 2013). The detected here urea, a product of uric acid mineralization, the affinity of the microbial community for N-acetyl-D-glucosamine utilization (chitin monomer) and high abundance of suspected chitin-degraders (*Vicinamibacteraceae* and *Chitinophagaceae* among others) indirectly corroborate this notion (Metze et al. 2024). Ammonia release from piscivorous bird-affected sites on the other hand seems to be rather rapid, as NH_4_^+^ concentration during the breeding season are often above 100 mg kg^-1^ of soil (Zwolicki et al., 2013; Zhu et al., 2012). The following nitrification step provides a direct substrate reservoir for microbial denitrification, a major N_2_O-releasing process (Yang et al., 2022). Nitrification usually is most active at neutral or slightly alkaline pH (Prosser, 2011). Soils of the OS site had an acidic pH, which has been observed before and seems to be a characteristic trait of the little auk affected nitrocoprophilous tundra soils (Zwolicki et al., 2013; Szymański et al., 2013). The high NO_3_^-^ concentrations observed by Zwolnicki et al. (2013) might be attributable to the transient alkalization of these soils following rapid ammonification of the labile guano components and NH_3_ release, allowing ammonia oxidizing bacteria to thrive. Those bacteria usually show affinity for high ammonia concentrations and display high nitrification rates (Rütting et al., 2021). Presence of bacterial genes coding for enzymes of the nitrification pathway was predicted based on the taxonomic diversity in this study, with members of the Nitrospirota and *Bradyrhizobiacea* as the most likely contributors (Daims et al., 2016; de Souza et al., 2013). Nitrification, N-acetyl-D-glucosamine degradation and processes related to ornithogenic soil formation lead to habitat acidification, thus potentially constricting further ammonia oxidation by bacteria (Breuning-Madsen et al., 2010; Poggere et al., 2016). Ammonia oxidizing archaea might take over the process as they appear to be abundant in Arctic tundra soil and prefer acidic, low NH_4_^+^ habitats (Siljanen et al., 2019). The resulting nitrate pool might be largely available to microbes as the nitrogen uptake by plants is highest at the start of the vegetation season with some of the arctic tundra plant species preferring other N forms like ammonia or organic N (free amino acids, urea) (Iversen et al., 2015). The presence of an anoxic niche beneath the vegetation cover, as suggested by the high relative abundance of butanol and acetate, likely hinders root penetration and nitrate uptake from deeper layers (Hädrich et al., 2012; Iversen et al., 2015). Consequently, these soil meet the major criteria for a complete denitrification to actively occur. The ratio of the predicted abundance of *nirK*+*nirS*/*nosZ* genes further supports this conclusion. However, several studies confirm, that low pH hampers the maturation process of the nosZ protein, which facilitates incomplete denitrification and consequently N_2_O emissions (Gaimster et al., 2018). Some argue, that such sites could host a complete denitrification process or even be a N_2_O sink, due to the presence of acid-tolerant denitrifiers, however this has yet to be evaluated for nitrocoprophilous tundra soils (Palmer and Horn, 2012). On that note, a few prominent bacterial families displayed a negative correlation with soil pH, hinting towards their acidophilic nature. One of them was the family *Gemmataceae* (Planctomycetota). Studies on denitrification-related gene abundance showed that permafrost-dwelling representatives of this group lack the *nosZ* gene, while still possessing NO-reductase genes (Frostegård et al., 2022). However, major players in N_2_O biosynthesis in these soils are likely to be the highly abundant members of the family *Vicinamibacteraceae*, which in a metagenomic study of Pessi et al. (2022) were recognized as possessing denitrification-related genes, including *nirK*, *norB* and *NorZ*. Our study shows how widespread these bacteria are in the soils of the Hornsund area, hinting at their remarkable adaptation capabilities. The relative abundance of this family has not correlated significantly with soil pH, so their preferred conditions remain elusive, as is their tendency for incomplete denitrification in acidic soils. However, the reduction of N_2_O to dinitrogen is the last reaction in the denitrification chain and also the least energy-yielding, suggesting that arctic tundra inhabiting *Vicinamibacteraceae* might have not evolved acid-resistant N_2_O reductases due to the unviability of this process in changing soil conditions (Koike and Hattori, 1975).

### 4.2 Biotic-related factors cause nitrate limitation in nitrogen-rich soils influenced by freshwater streams

The second habitat, that according to our data could have the potential to emit N_2_O in considerable amounts was the ST site. Soils of this area are influenced by the streams of the Fuglebekken catchment, which transport dissolved substances from the ornithocoprophilous tundra (site OS) to the Hornsund fjord (Szymański et al., 2016). Therefore, the nutrients at this site largely originate from the little auk nesting site upstream, resulting in many of the geomicrobiological features to be shared between those sampling points. However, several key characteristics of N_2_O-emitting hotspots are missing. Despite the presence of a large N-pool and a low C:N ratio, nitrates were not detected in these soils, although ammonia showed the highest concentration score among investigated samples. This is somewhat consistent with the findings of Zwolicki et al. (2013), where NO_3_^-^ levels sharply decline with increasing distance from the colony, whereas NH_4_^+^ is detected beyond the colonies immediate vicinity. This could be attributed to the volatile nature of ammonia (Otero et al., 2018). However, relatively high NH_4_^+^ concentration, neutral pH and diverse (phylogenetically and metabolically) bacterial community in ST soils should promote nitrification by the highly active ammonia-oxidizing bacteria (Rütting et al., 2021). Thus, nitrates at this site are either not produced or rapidly sequestered by plants and microorganisms. An argument for the former could be the high water saturation of the examined soil, which is often the cause of anoxic conditions, unfavorable for nitrifying microbes (Voigt et al., 2020). Interestingly, the relative abundance of the phylum Nitrospirota was the lowest at this site, although the abundance of nitrification-involved genes was quite high, which supports the latter hypothesis on nitrate deficit. Consequently, whatever amount of nitrates is produced in these conditions, it might get taken up by diverse metabolic processes, including primary production (Fowler et al., 2013). Several features suggest an active primary production process at this site, in contrast to the ornithogenic tundra soils, where the degradation of guano and other bird-derived materials was the main nutrient-acquisition strategy (Skrzypek et al., 2015). In ST soil there were detectable amounts of highly labile compounds like glycerol or uracil, which if not actively released by living cells, remain undetectable as they degrade rapidly (Ferrer-González et al., 2023). The high amount of aliphatic compounds with chains of varying length also suggests anabolic activity indicative of cell membrane lipid synthesis or microbial storage materials (Zelles, 1999; Wältermann et al., 2007). EcoPlate responses showed that the ST microbial community was able to efficiently metabolize plant-derived nutrients like mannitol or cellobiose but also galacturonic acid, which is a monomer of pectin, suggesting a development of a diverse rhizospheric community dependent on plant litter and plant exudates (Znój et al., 2022). The relatively high concentrations of biogenic metals at this site might also be explained by the combined activity of growing plants and their accompanying microflora due to siderophore production and other means of bioleaching (Timofeeva et al., 2022). Nitrates can also be used in dissimilatory reduction resulting in ammonia release, a process conducted *inter alia* by the bacteria of the phylum Chloroflexi (West-Roberts et al., 2021). The relative abundance of members of this phylum was the highest in ST soils, where their metabolic activity could explain the lack of nitrates but also the high ammonia concentrations. Notably, dissimilatory nitrate reduction generates N_2_O as a byproduct, although to a lesser extent than the incomplete denitrification process (Torres et al., 2021). The ST bacterial community, as compared to the OS area, displays the highest *nirK*+*nirS*/*nosZ* ratio of any examined soil community, which makes it, at least microbiologically, prone do emit N_2_O during denitrification (Voigt et al., 2020). That the majority of arctic tundra denitrifies have truncated denitrification pathways was reported before, suggesting that the vast stretches of the tundra biome are a prolific nitrous oxide emitter (Pessi et al., 2022). However, as in the case of the ST site, biotic-related factors are likely to restrict nitrate availability, thus minimizing the threat of possible mass N_2_O release (Voigt et al., 2020).

### 4.3 High Arctic soils without external nutrient inputs remain N poor and show traits of a N_2_O sink

Soils of the remaining sites displayed significantly lower total nitrogen concentrations compared to the OS and ST site, highlighting the impact of the *Alle alle* colony. The moss-covered MC site could have some capacity for N_2_O emissions given that total N amounts were relatively high in these soils with a detectable level of NH_4_ ^+^. However, considering that the majority of carbon at this site was bound by recalcitrant organic matter (long chain alkanes and phenolic compounds), the nitrogen was likely also immobilized in organic debris and/or living cell biomass, making it unavailable for N_2_O-producing microbial processes (Vasilevich et al., 2018). Furthermore, labile nitrogen could be produced and utilized only within the narrow constrains of the moss rhizosphere as there is evidence for an active nitrification and dinitrogen fixation promoted by moss communities, making those nitrogen forms less available to other, non-symbiotic soil microbiota (Hayashi et al., 2016; Rousk et al., 2017; Ramond et al., 2022). Additionally, moss coverage is known for delaying the thaw of the organic permafrost layer, while also reducing the moisture of the underlying soil, creating sub-optimal conditions for anaerobic N_2_O emitters (Gornall et al., 2007). There are reports of mosses as N_2_O sources, however these findings have yet to be verified *in situ* for Arctic moss species (Lenhart et al., 2015). The sites HG, HS and PG displayed the lowest total nitrogen concentrations and no detectable NO_3_^-^ and NH_4_^+^ amounts, making them least likely to be N_2_O emission hotspots. Some of those sites could rather act as a N_2_O sink (“coldspot”), especially the post-glacial site within the Hans Glacier moraine system. It displayed several traits indicative of potential nitrous oxide sink, most notably a high C:N ratio and low *nirK*+*nirS*/*nosZ* ratio. The high pH of the soil is also beneficial, especially due to its high buffering capacity as suggested by the high calcium concentration (Kabala and Zapart, 2012). Low soil moisture found at this site may contradict the notion of it being a N_2_O “coldspot”, however this is a highly labile factor, strongly dependent on seasonal changes and current weather conditions. Furthermore, the relatively high C content found in those soils could be stimulating respiration, leading to anaerobic micro niche development, where denitrification could take place (Chapuis-Lardy et al., 2007). Glacier forelands, if confirmed as nitrous oxide sinks, have the capacity to significantly lower atmospheric N_2_O levels as their surface area is rapidly increasing due to the global temperature rise and glacier retreat (Hugonnet et al., 2021).

## 5. Conclusions and Future Directions

Our study identifies little auk (*Alle alle*) breeding colonies as probable hotspots of nitrous oxide (NLO) emissions in Arctic tundra soils. The ornithogenic soils at the OS site displayed several hallmarks of microbial NLO production, including elevated nitrogen concentrations, the presence of nitrates, an abundant and functionally diverse microbial community and strong probability of incomplete denitrification under acidic conditions. The persistence of relatively high ammonium concentrations past the breeding season suggests a unique, planktivorous bird–associated mechanism of gradual nitrogen release linked to the slow decomposition of chitin-rich guano. This sets little auk colonies apart from piscivorous seabird colonies, where nitrogen turnover is more rapid.

In contrast, the ST site, though nutrient-rich and microbiologically prone to denitrification, appears limited by nitrate availability. Biotic processes such as plant uptake, microbial assimilation, and dissimilatory nitrate reduction likely suppress large-scale NLO emissions, despite high ammonium levels and metabolically versatile microbial communities. Other sites lacking ornithogenic input showed low nitrogen availability, with some, particularly the Hans Glacier foreland, potentially functioning as NLO sinks due to favorable soil chemical and microbial traits.

Taken together, these findings suggest that nutrient subsidies from planktivorous seabirds play a critical but underappreciated role in shaping Arctic greenhouse gas dynamics, generating localized hotspots of NLO emissions amidst generally nitrogen-poor soils.

Future work should focus on:

- Quantifying temporal variability and “hot moments” of NLO fluxes across the seabird breeding cycle.
- Experimentally testing the role of chitin and other guano-derived compounds in regulating ammonium release and microbial activity.
- Elucidating the contributions of acidophilic and acid-tolerant denitrifiers, particularly *Vicinamibacteraceae*, to incomplete denitrification in ornithogenic soils.
- Assessing the long-term role of glacier forelands as potential NLO sinks in the context of rapid Arctic deglaciation.
- Integrating microbial functional data with direct NLO flux measurements to refine ecosystem-scale greenhouse gas budgets.

By combining microbial ecology with biogeochemical monitoring, future studies will help clarify the balance between emission hotspots and sinks in Arctic tundra, ultimately improving predictions of how climate change and shifting seabird populations may alter NLO dynamics in polar regions.

## CRediT authorship contribution statement

**Julia Brzykcy:** Conceptualization; Methodology, Funding acquisition, Investigation, Formal analysis, Resources, Data Curation, Validation, Visualization, Writing - Original Draft, Writing - Review & Editing.

**Renata Matlakowska:** Conceptualization; Methodology, Supervision, Funding acquisition, Project administration, Writing – Review & Editing.

**Jakub Grzesiak:** Methodology, Software, Formal analysis, Resources, Data Curation, Writing – Review & Editing, Writing – Original draft, Visualization.

## Funding

This work was supported by:

– The program of the Ministry of Science and Higher Education (Poland) - Excellence Initiative - Research University (2020-2026) No. BOB-IDUB-622-858/2023;
– Priority Research Areas - Science for the Planet and Polar Mission 2021, Edu Arctic (Institute of Geophysics Polish Academy of Sciences).

## Acknowledgements

The authors express their gratitude to Robert Stasiuk and Bartosz Rewerski (Laboratory of Instrumental Environmental Analysis; Department of Biology, University of Warsaw) for their cooperation in the geochemical analyses, as well as organizers of the Polar Mission 2021, Edu Arctic project (Institute of Geophysics Polish Academy of Sciences) and members of the 44^th^ Polar Expedition of the IG PAS to Polish Polar Station Hornsund for their support during fieldwork.

## Data availability

Illumina reads were deposited in the NCBI Sequence Read Archive (SRA) as BioProject PRJNA1224745. Other data will be made available on request.

